# Leaf economics explained by optimality principles

**DOI:** 10.1101/2021.02.07.430028

**Authors:** Han Wang, I. Colin Prentice, Ian J. Wright, Shengchao Qiao, Xiangtao Xu, Kihachiro Kikuzawa, Nils Chr. Stenseth

## Abstract

The worldwide leaf economics spectrum relates leaf lifespan (LL) to leaf dry mass per unit area (LMA)^1^. By combining three well-supported principles^2-4^, we show that an isometric relationship between these two quantities maximizes the leaf’s net carbon gain. This theory predicts a spectrum of equally competent LMA-LL combinations in any given environment, and how their optimal ratio varies across environments. By analysing two large, independent leaf-trait datasets for woody species^1,5^, we provide quantitative empirical support for the predicted dependencies of LL on LMA and environment in evergreen plants, and for the distinct predicted dependencies of LMA on light, temperature, growing-season length and aridity in evergreen and deciduous plants. We thereby resolve the long-standing question of why deciduous LMA tends to increase (with increasing LL) towards the equator, while evergreen LMA and LL decrease^6^. We also show how the statistical distribution of LMA within communities can be modelled as an outcome of environmental selection on the global pool of species with diverse values of LMA and LL.

## Main

Plants are subject to powerful selective forces in every aspect of their lives, favouring economic efficiency of vegetative processes as a necessary condition of reproductive success in a competitive milieu. We will show here how the worldwide leaf economics spectrum (LES) – the generalization that thicker and/or denser leaves (high LMA) have longer lifespans (high LL)^1^ – arises as a consequence of leaf-level maximization of net carbon gain.

LL and LMA vary several hundredfold across vascular plants, and up to tenfold among co-occurring species^1,7^. Central values of LL, LMA and their ratio also shift along climatic gradients^8-10^, but the reasons are not well understood. The environmental controls on LL and LMA remain unclear, as do the different responses of deciduous and evergreen species^11^. In particular, why do the leaves of deciduous trees and shrubs tend to become thinner (as their lifespan declines) poleward, while those of evergreen species become thicker^12^ and longer-lived^13^? Better understanding of the LES is required: not least for more robust modelling of the global carbon cycle and its interactions with climate, as LL and LMA directly influence carbon cycling in terrestrial ecosystems by determining the carbon requirements for leaf construction and turnover^14-16^.

Recent research has shown that eco-evolutionary optimality hypotheses can predict both known and unexpected patterns of relationship among plant traits, and between traits and environment^17^. The increasing availability of large plant trait data sets allows quantitative assessment of such predictions^18^. Much research has focused on leaf physiological traits, like photosynthetic capacity, that can change within the same plants (and even the same leaves) in response to seasonal environmental cues^19,20^. Unlike physiological traits, however, LMA and LL generally show more muted intraspecific responses to temporal changes and geographic gradients in environmental factors. Differences in the phenotypic plasticity and/or genotypic adaptability of traits are revealed by comparison of within-species and between-community trends along environmental gradients^21^. In a continental-scale analysis, intraspecific shifts in LMA showed only about one-third of the response to environmental variation shown by community-mean LMA^21^, compared to (for example) > 90% for the ratio of leaf-internal to ambient CO_2_, a key physiological trait involved in photosynthesis^22^. Nonetheless, the LES can be demonstrated across species worldwide^1^ and the associated traits shift with environment in consistent ways, strongly suggesting the existence of a common organizing principle^8-10,23^.

### An optimality theory for the leaf economic spectrum

An eco-evolutionary optimality approach is developed here to account for the environmental controls on leaf economics for woody species in a quantitative framework. First, we consider the relationship between LMA and LL that maximizes the leaf life-cycle average net carbon gain, i.e. net photosynthesis minus the associated (amortized) tissue construction costs^2^. This relationship depends on the rate at which photosynthesis declines with leaf age^3^. Second, the ageing rate increases in proportion to the leaf’s initial photosynthetic capacity, as measured by the maximum rate of carboxylation at a standard temperature of 25°C (*V*_cmax25_), and decreases in proportion to LMA^3^. Third, photosynthetic capacity itself is optimized to the physical environment, following the coordination hypothesis^4,24,25^. Combining these three principles (see Methods for derivations) we obtained the following general relationship between LMA and optimal LL (LL_ev_) in evergreen leaves:

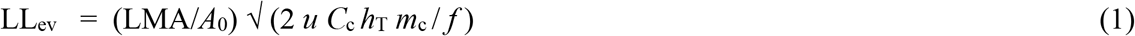

where *A*_0_ is the leaf’s initial photosynthetic rate; *f* is the growing-season length, as a fraction of the year; *u* is a constant that relates the ageing rate to LMA and photosynthetic capacity; *C*_c_ is a multiplier for the total carbon costs, including required investments in non-leaf tissues, associated with leaf construction; *h*_T_ is the Arrhenius function relating *V*_cmax_ to temperature (*T*), equal to unity when *T* = 298.15 K; and *m*_c_ is the ratio of the Rubisco-limited photosynthetic rate to *V*_cmax_, which is a function of the intercellular CO_2_ partial pressure and the affinity of Rubisco for CO_2_ versus O_2_. The coordination hypothesis leads to the prediction that *A*_0_ is proportional to the photosynthetic photon flux density (PPFD) absorbed by the leaf^20^.

This theory predicts the existence of a spectrum of values of both quantities, from low LMA and LL to high LMA and LL, resulting in equal rates of net carbon gain (see Methods). The optimal LL in a given environment is proportional to LMA, and inversely proportional to absorbed PPFD and the square root of growing-season length. We show here (Figure 1a-c, Table S1) that these predictions are similar to the corresponding partial effects independently inferred from a multiple regression fitted to the measurements on evergreen species in the GlopNet data set^1^, which provides the largest available worldwide compilation of paired data on LMA and LL. The regression for evergreen species captures 42% of the variation in LL. Evergreen LL is proportional to LMA (log-log slope ≈ 1) as predicted. The relationship of LL to PPFD is negative as predicted, and the relationship of the LL residuals to growing-season length is also negative (*P* < 0.1) as predicted, with predicted slopes similar in magnitude to fitted values (although falling just outside their predicted confidence intervals).

**Fig. 1.**
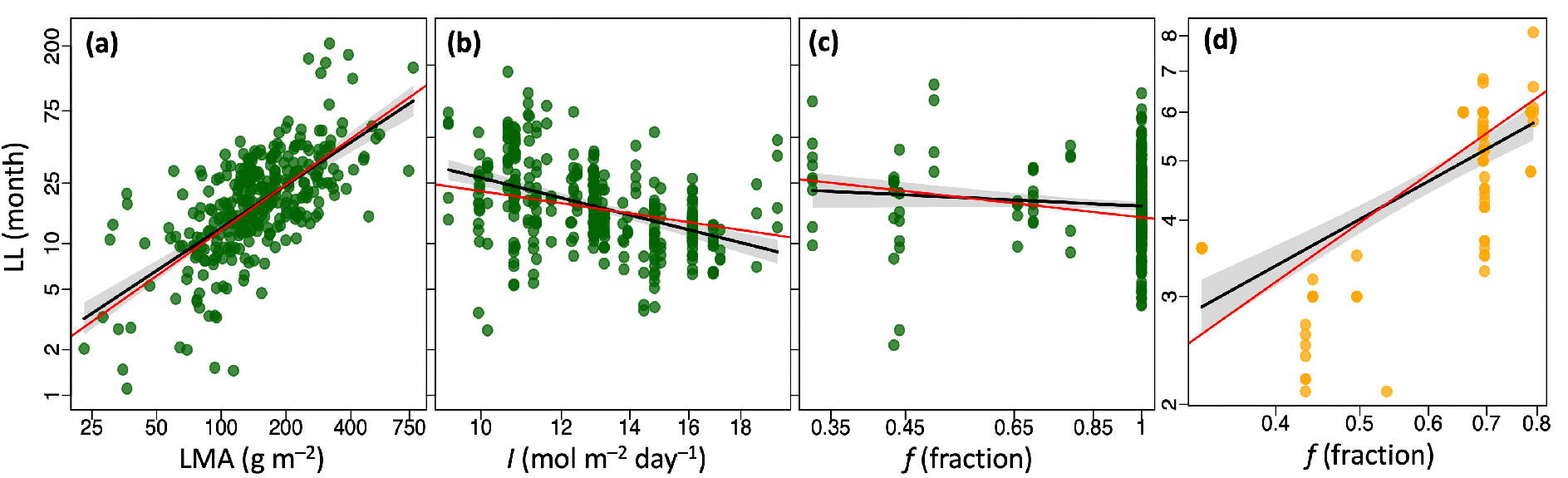
Partial residual plots from the regression of observed leaf longevity (LL) against explanatory variables for evergreen and deciduous species. **a**, LMA: leaf mass per area. **b**, *I*: site-mean leaf absorbed Photosynthetic Photon Flux Density. **c and d**, *f*: growing-season length as a fraction of the year. Predicted relationships in red, fitted in black with 95% confidence interval in grey; evergreen species in green, deciduous in orange. All axes are on log scales. Data from the GlopNet trait data set^1^.

Due to the different behaviour of evergreen and deciduous species in the unfavourable season, the same optimality framework leads to a different equation that describes net carbon-gain maximization in deciduous species (see Methods for derivation):

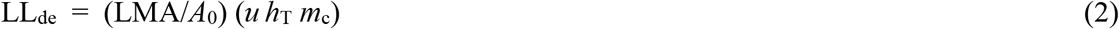

In addition to this optimality criterion, LL_de_ is tightly constrained by growing-season length. Among the deciduous species in GlopNet, growing-season length is indeed nearly proportional to LL (0.79 ± 0.08, 1 s.e.), and explains about half of the observed variation in LL (Figure 1d, Table S1).

Equations (1) and (2) imply that for any given LL, both evergreen and deciduous LMA should increase in direct proportion to absorbed PPFD. All else equal, LMA should also decrease by 2.6% K^−1^ with increasing temperature in evergreen species, and at about twice that rate in deciduous species (see Methods). The response of LMA to growth temperature is mainly a consequence of the thermal acclimation (down-regulation) of carboxylation capacity, which implies slower leaf ageing at higher temperatures, and therefore reduces the carbon investment needed to produce a unit of photosynthate. High LMA is also expected (and found) in plants of arid environments, because maintaining turgor at low leaf water potential requires leaves to be stiff^26^, and because thin leaves have insufficient heat capacity to prevent overheating during transient low-wind episodes^27^.

### The environmental dependencies of leaf mass per area

This theory predicts different environmental dependencies of LMA in evergreen and deciduous plants. To quantify these dependencies, we must consider how environmental constraints act on prior probability distributions of LMA and LL. The principle is illustrated in Figure S2, where straight lines in the log-log plot of LL versus LMA – representing the optimal LES for two sites in different environments – intersect lognormal distributions of both variables. In deciduous plants, site-mean LL is constrained by growing-season length. In evergreen plants, site-mean LMA is predicted at the mean of the distribution formed by the intersection of the lines with the prior distribution. The resulting predicted sensitivities of evergreen LMA to temperature and PPFD are only half those for deciduous LMA (see Methods). For growing-season length, they are only a quarter of those for deciduous LMA.

These predictions are fully supported by analysis of data in the China Plant Trait Database (CPTD)^5^, which provides a larger LMA data set than GlopNet for temperate and boreal deciduous species, many more observations for deciduous species in particular, and more than three times as many sites. Growing-season PPFD, temperature and an index of plant-available moisture (see Methods) all influence the LMA of evergreen species (Figure 2a-c, Table S2).

**Fig. 2.**
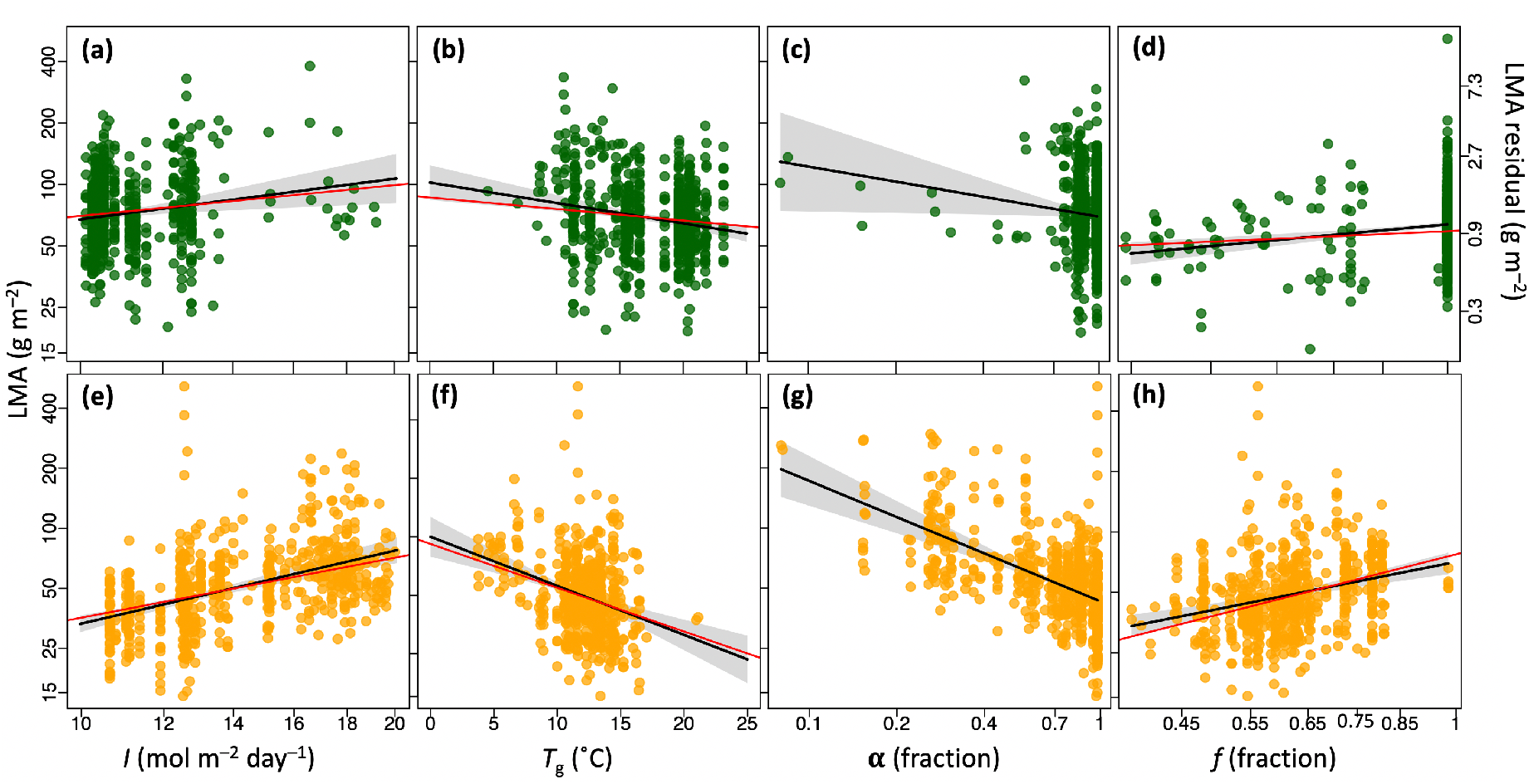
Partial residual plots from the regression of observed leaf mass per area (LMA) against environmental variables for evergreen and deciduous species. **a and e**, *I*: site-mean leaf absorbed Photosynthetic Photon Flux Density. **b and f**, *T*_g_: the mean temperature of the growing season, when daily temperature is above 0°C. **c and g**, *α*: moisture index (the ratio of actual to equilibrium evapotranspiration). **d and h**, *f*: growing-season length as a fraction of the year. Predicted relationships in red, fitted in black with 95% confidence interval in grey; evergreen species in green, deciduous in orange. All axes except for *T*_g_ are on log scales. Data from an extended version of the China Plant Trait database^5,28^.

The predictive power of this relationship is limited (13%), however, presumably because environmental predictors alone can only explain a limited proportion of the variation in LMA (additional variation being driven by variation in LL that is independent of environment) because the species span a wide range of LL. Growing-season length is also a significant predictor of the residual variation in evergreen LMA (Figure 2d, Table S2). The same four environmental predictors account for more than half of the variation in LMA among deciduous species in the CPTD (Figure 2e-h, Table S2), where growing-season length is expected to largely determine LL and so to be proportional to LMA, as observed. The confidence intervals of the fitted slope coefficients for PPFD, temperature and growing-season length all include the predicted values for both evergreen and deciduous species.

For evergreen species, whose LL is not directly constrained by the growing-season length, it is useful to predict not only a community-mean value for LMA but also a statistical distribution representing the site-specific LES. We fitted independent lognormal distributions of LMA and LL based on GlopNet data, and imposed equation (1) as a constraint. This method successfully predicted within-community LMA distributions in evergreen temperate forests, tropical rainforests and woodlands (Figure 3). The community-mean LMA values of temperate forests and tropical rainforests are close to the global mean value of LMA derived from the GlopNet data set. They are predicted to be similar because growth temperature and growing-season PPFD and duration are positively correlated, but to have opposite effects on LMA. The community-mean LMA of woodlands (Figs 3e,f) however shifts far from the global mean, because of the additional ecophysiological constraint imposed by aridity.

**Fig. 3.**
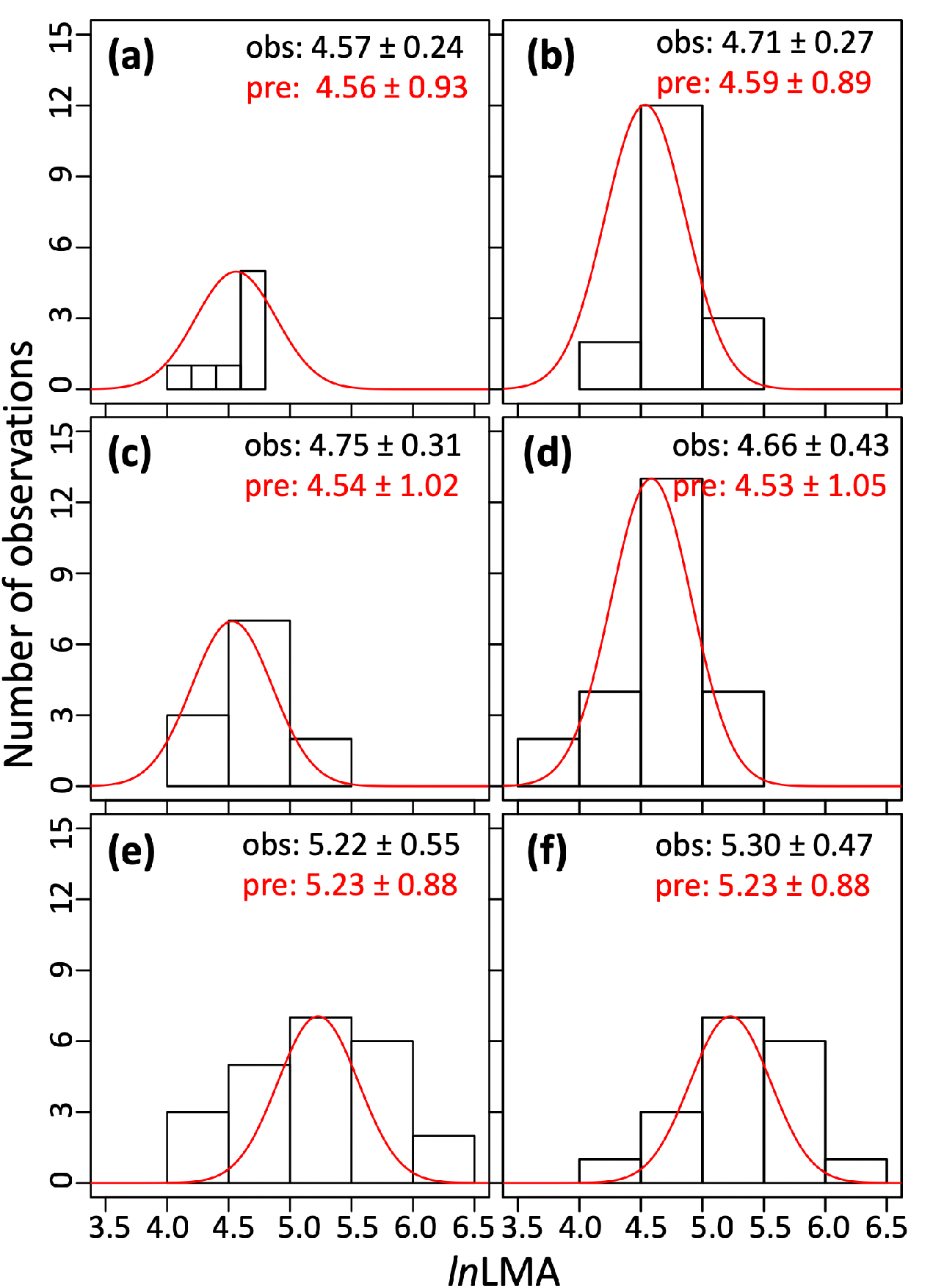
Observed and predicted within-site distributions of leaf mass per area (LMA). **a and b**, temperate forests. **c and d**, tropical rainforests. **e and f**, woodlands^29^. Means and standard errors of the observed (obs) and predicted (pre) distributions are shown. Predicted probability distribution functions (red curve) are scaled in height to match the observed frequency distribution. Data from the GLOPNET trait data set ^1^. Sites with the largest sample size are selected; see detailed site information in Supplementary Table S3.

This theory accounts for the contrasting latitudinal patterns in LMA between deciduous and evergreen species^6^, as shown in Figures 4 and S3. The expected effect of increasing temperature towards the equator on LMA is negative for both leaf types. However, in deciduous species, this influence is outweighed by the effects of increasing growing-season PPFD and length, both of which favour increased LMA.

**Fig. 4.**
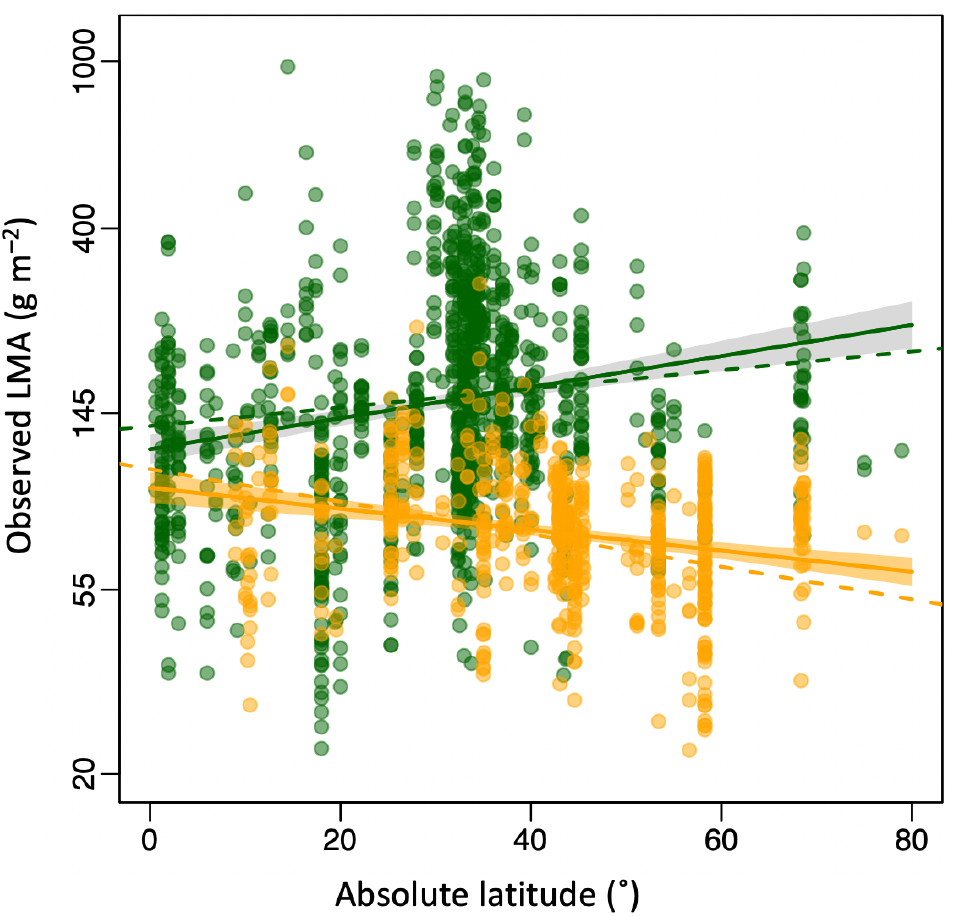
Observed and predicted trends of leaf mass per area (LMA) from tropical to polar regions. Data and fitted regression lines with 95% confidence intervals are in green for evergreen and orange for deciduous species. Dashed lines are theoretical predictions. Data from the GlopNet trait data set^1^.

Carbon-balance optimality provides a quantitative explanation for the existence of the LES, and for its systematic variations with environmental conditions. The resulting patterns are key to understanding the biogeographical dimension of leaf economics. Unlike variations in physiological traits, these patterns are primarily determined by environmental selection among species rather than plasticity and genetic adaptation within species^22,30^. However, plants do have some plasticity for LMA, as has been shown experimentally^31^. Experimental effects on LMA include positive responses to PPFD, CO_2_ and aridity and a negative response to growth temperature^31,32^ – all responses in the same direction as predicted here, but generally smaller in magnitude.

Plants have been subject to profound changes in climate during their evolutionary history. Large, and sometimes rapid, changes in recent geological times have resulted in repeated large-scale reassortments of species and consequent major changes in community composition^33,34^. Current anthropogenic environmental changes are adding to the forces shaping plant communities, in ways that remain only partly understood^16^. Optimality theory may serve not only to solve the puzzles of global phytogeography, but also to move plant functional ecology from empirical description to robust theoretical understanding^17^.

## Methods

### An optimality framework for leaf economics

The optimality model of leaf longevity (LL, day) proposed by Kikuzawa^2^ rests on two assumptions. First, daily photosynthesis declines linearly from an initial maximum (denoted by *A*_0_, g biomass m^−2^ day^−1^) with increasing age. Second, LL maximizes the leaf’s lifetime-average net carbon gain (accumulated daily photosynthesis averaged over LL, minus the initial structural investment amortized over LL). Building on Kikuzawa’s original work, we propose a new, unified optimality model for both deciduous and evergreen species. For deciduous species, we extend the period for maximizing average net carbon gain from the leaf life span to the whole leaf life cycle (LC, day), i.e. the period from the formation of a leaf until its replacement by a new one. LC is thus equal to LL for evergreen species, but to 365 days for deciduous species. We further simplify the derivation by disregarding leaf maintenance respiration, as it is relatively small, and nearly proportional to photosynthetic capacity^35^. The daily net carbon gain (*g*) is then:

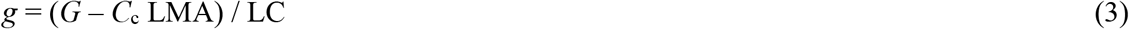

where *G* (g biomass m^−2^) is the accumulated photosynthetic carbon gain over the life cycle. The initial structural investments are represented by the product of LMA and *C*_c_, where LMA is leaf mass per area (g biomass m^−2^), and *C*_c_ (> 1, gC gC^−1^) is the total construction cost, including non-leaf tissues, of a unit of leaf mass. *G* is represented differently for deciduous and evergreen species, since deciduous leaves can carry out photosynthesis during their whole life span, whereas evergreen leaves are inactive during the non-growing season.

We use subscripts ‘ev’ and ‘de’ to distinguish evergreen and deciduous species. Thus for evergreen plants:

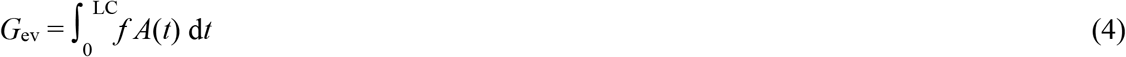

where *A*(*t*) = *A*_0_ (1 – *t*/*b)* is the assimilation rate (g biomass m^−2^ day^−1^) as a function of age (*t*, days) and *b* (days) is the leaf age at which carbon assimilation would decline to zero. *f* is the proportion of the year that is favourable for growth (day day^−1^). Since LC_ev_ = LL_ev_, the integral in (4) is equal to *f A*_0_ [LL_ev_ – (LL_ev_)^2^/2*b*], and equation (3) collapses to Kikuzawa’s model^2^:

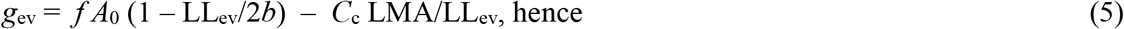

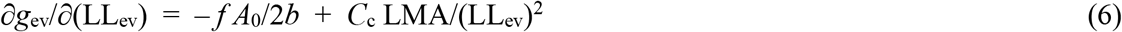

Setting equation (6) to zero yields an expression for optimal LL_ev_ (days) in evergreen plants:

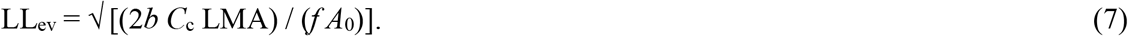

For deciduous plants:

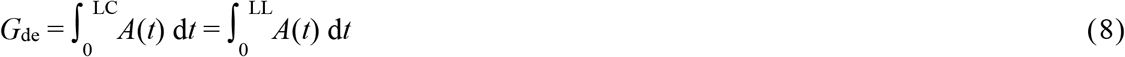

The integral in equation (8) equals *A*_0_ [LL_de_ – (LL_de_)^2^/2*b*] and thus

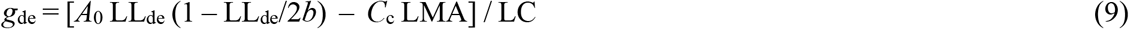

In deciduous species LC = 365, thus

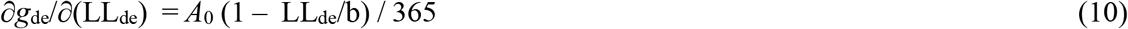

Setting equation (10) to zero yields a simple equation for optimal LL (days) in deciduous plants:

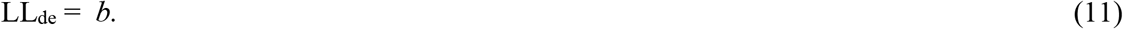

We implemented the model of *b* proposed by Xu *et al*.^3^ in the framework described above. Xu *et al*. demonstrated that *b* has a positive relationship with LMA, and a negative relationship to the maximum capacity of carboxylation at 25°C (*V*_cmax25_, μmol C m^−2^ s^−1^), with scaling coefficients close to 1 and –1 respectively. Therefore, *b* can be expressed as:

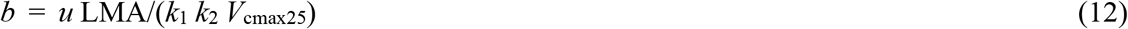

Here *u* ≈ 768 ± 71 (dimensionless), estimated from a meta-analysis of data on 49 species across temperate and tropical biomes^3^. The scaling factors are *k*_1_ = 30 g biomass mol C^−1^ and *k*_2_ = 0.0864 s mol C day^−1^ μmol^−1^ C.

### The optimality model of leaf photosynthesis

The coordination hypothesis, supported by extensive field and experimental evidence^25,35^, states that the maximum capacity of carboxylation at growth temperature (*V*_cmax.gt_, μmol C m^−2^s^−1^) acclimates to the daytime environment so that the Rubisco-limited photosynthetic rate (*A*_C_, μmol C m^−2^ s^−1^) tends to equality with the electron transport-limited rate (*A*_J_, μmol C m^−2^ s^−1^). In other words, *A*_C_ = *A*_J_. This acclimation is optimal because it avoids investment in excess photosynthetic capacity, while allowing full use of the available light^4,24^. Combined with the standard biochemical model of photosynthesis^36^, the coordination hypothesis predicts *V*_cmax.gt_:

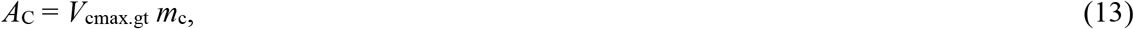

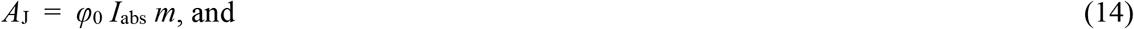

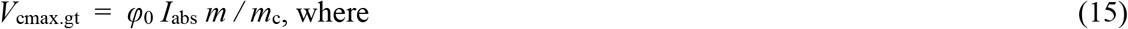

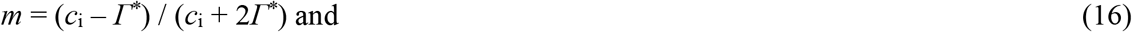

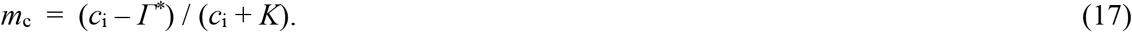

Here, *φ*_0_ is the intrinsic quantum efficiency of photosynthesis (μmol C μmol^−1^ photon), which we assume to depend on temperature as described in ref. ^37^. *I*_abs_ is the leaf-absorbed PPFD (μmol photon m^−2^ s^−1^) to which *V*_cmax.gt_ acclimates, *m*_c_ and *m* are the CO_2_ limitation terms for *A*_C_ and *A*_J_ respectively, *c*_i_ is the leaf-internal partial pressure of CO_2_ (Pa), *Γ*^*\**^ is the photorespiratory compensation point (Pa), and *K* is the effective Michaelis-Menten coefficient of Rubisco (Pa). Optimal stomatal regulation, according to the least-cost hypothesis^38^, yields the following relationship of *c*_i_ to environmental variables, including vapour pressure deficit (*D*, Pa) and *c*_a_:

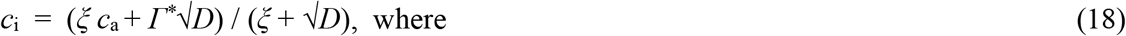

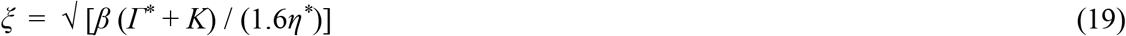

The dimensionless parameter *β* in equation (19) is the ratio of unit costs for the maintenance of carboxylation and water transport capacities, evaluated at 25°C. The term *η*^*\**^ is the viscosity of water (normalized by its value at 25°C), which declines with increasing temperature, thereby reducing the cost of water transport and thereby favouring higher *c*_i_. The value of *c*_i_ thus depends on temperature via *η*^*\**^, *Γ*^*\**^ and *K*, as well as on *D* and *c*_a38_. Analysis of a global leaf stable carbon isotope dataset^20^ has provided empirical support for the separate dependencies of the *c*_i_:*c*_a_ ratio on temperature, vapour pressure deficit and elevation implied by equations (18) and (19), and an estimated value of *β* ≈ 146 (refs ^20,39^).

The temperature dependencies of *Γ*^*\**^, *K* and *V*_cmax_ within normal physological ranges can be described by the Arrhenius function^40^ with different activation energies (the formulation for *K* is somewhat more complex as it depends on the affinities of Rubisco for CO_2_ and O_2_ and their distinct activation energies, while both *Γ*^*\**^ and *K* are influenced by atmospheric pressure: see ref. ^40^ for details). *V*_cmax25_ is related to *V*_cmax.gt_ by:

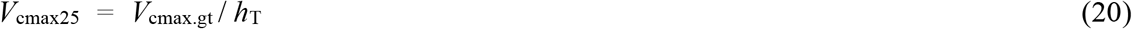

where *h*_T_ is the Arrhenius function for *V*_cmax_.

We further assume that the coordination hypothesis defines the initial photosynthetic capacity of the leaf. Thus,

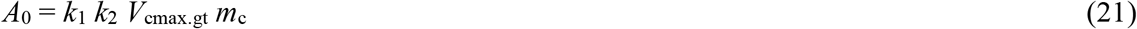

Here, *k*_1_ and *k*_2_ together convert the unit of *V*_cmax.gt_ from μmol C m^−2^ s^−1^ into g biomass m^−2^ day^−1^ to match the units of *A*_0_.

### Optimal leaf longevity in evergreen species

Substituting equations (12)–(21) into (7) yields equation (1). Substituting equation (7) into (5) also yields an expression for an evergreen leaf’s average daily net carbon gain:

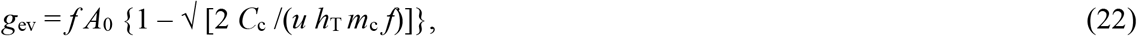

which is independent of either LMA or LL. In other words, any combination of LMA and LL that satisfies the optimality criterion will yield the same daily net carbon gain.

The coordination hypothesis also relates the initial rate of carbon assimilation to the electron-transport limited photosynthetic rate:

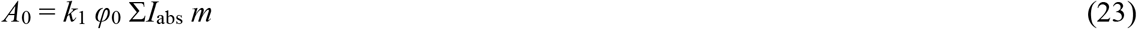

where Σ*I*_abs_ is the integrated instantaneous *I*_abs_ through a day (mol photon m^−2^ day^−1^). It follows that:

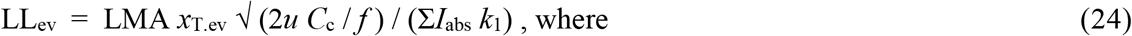

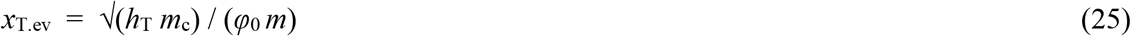

The composite term *x*_T.ev_ includes the temperature-dependent variables. We express the parameter *C*_c_ as a function of LL, LMA and environmental predictors by rearranging equations (24) and (25) as follows:

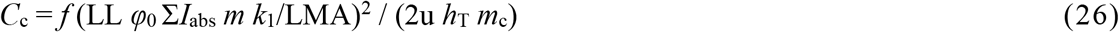

Based on all paired observations of LL and LMA in evergreen species in GlopNet (326 samples), we estimated a median value of *C*_*c*_ = 23 ± 5.12 gC gC^−1^ (mean ± s.e.).

### Optimality leaf longevity in deciduous species

Substituting equations (12), (15) and (20) into (11) yields equation (2). Substituting equation (2) into (9) also yields an expression for a deciduous leaf’s average daily net carbon gain:

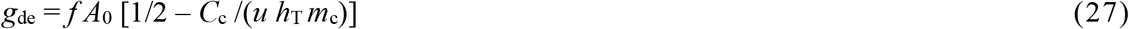

As in evergreen species, *g*_de_ is independent of either LMA or LL. The different formulae for *g*_de_ and *g*_ev_ potentially allow modelling of competition between the two strategies.

Following similar logic to that applied to evergreens, equation (23) is then substituted into equation (2), leading to a prediction of LL_de_ from LMA and environmental variables:

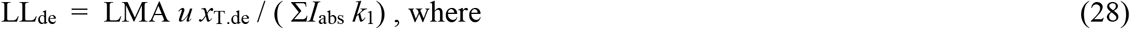

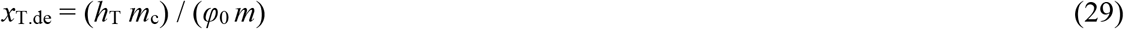

#### Linearized models for optimal LL

To facilitate comparisons of data with theoretical predictions, equation (24) can be linearized:

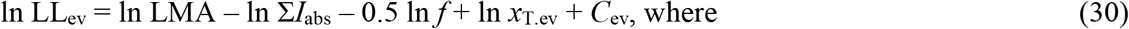

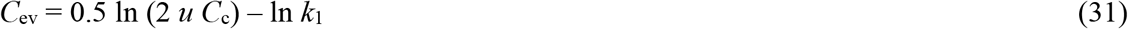

By taking the partial derivative of ln *x*_T.ev_ with respect to temperature, we calculated that LL_opt.ev_ increases with growth temperature (*T*_g_, K) by 2.6% K^−1^ under standard environmental conditions. Thus, ln *x*_T.ev_ can be replaced by the sum of 0.026 *T*_g_ and ln *x*T._ev_ at the reference condition (ln *x*_T0.ev_). *D* and *z* are also expected to influence LL_opt_ slightly via their effects on optimal *c*_i_, but these effects are too small to be detected.

Similar logic yields a linearized model of LL for deciduous species from equation (28):

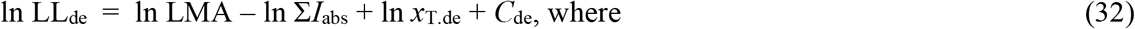

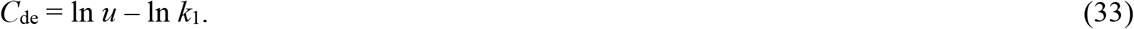

The thermal sensitivity of LL_de_ is twice of that in evergreen species. Thus, ln *x*_T.de_ can be replaced by 0.052 *T*_g_ plus its reference value (ln *x*_T0.de_).

#### Distributions of evergreen LMA and LL

Different combinations of LMA and LL consistent with the optimality criterion result in equal net carbon gain, implying the existence of a LES along which species with different LMA are equally competitive. Nonetheless, within-site distributions of LMA and LL are known to vary systematically with climate. To account for this variation, it is necessary first to take account of their prior distributions in the species pool. We fitted lognormal distributions to observed LMA and LL data independently in the full GlopNet dataset (Figure S1). Equation (24) was then imposed as a constraint, representing environmental filtering of the species pool. This procedure generates site-specific lognormal distributions, which serve as predictions of the probability density functions of LMA and LL subject to the optimality criterion as illustrated in Figure S2.

#### Environmental dependencies of LMA for evergreen species

Equation (30) can be re-configured as an expression for the optimal LMA of evergreen species:

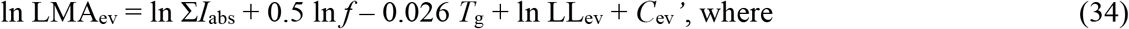

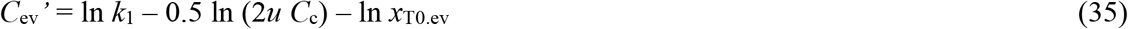

Optimal LMA depends on LL as well as on environmental factors. For a given LL, equation (34) implies that LMA (all else equal) should increase in proportion to absorbed PPFD and the square root of growing-season length, but should decrease with temperature by 2.6% K^−1^. The prior normal distributions of ln LMA and ln LL, together with their predicted proportional relationship, allow LMA to be predicted from environment alone (Figure S2). Here, we consider Σ*I*_abs_ as an example to make the derivation, noting that the effects of the other environmental variables (*f* and *T*_g_) can be derived following the same logic. We specify equations (34) and (35) at two sites with different light conditions (Σ*I*_1_ and Σ*I*_2_) but the same growing-season length and growth temperature. The subscripts ‘1’ and ‘2’ in the equations below distinguish the two sites:

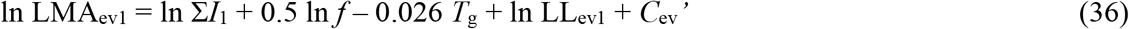

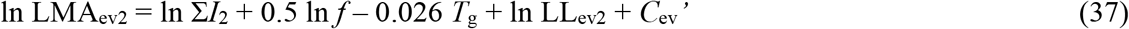

Thus,

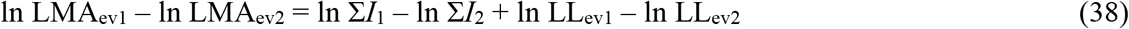

The normal distribution constraint induces a line perpendicular to the optimality lines for evergreen species (Figure S2):

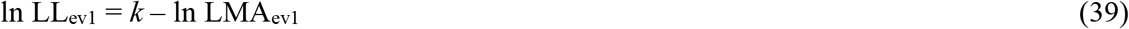

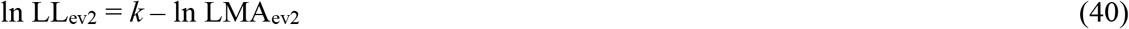

where k is the intercept of this line. Therefore,

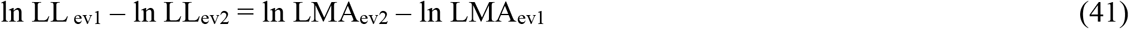

Combining equations (38) and (41) yields:

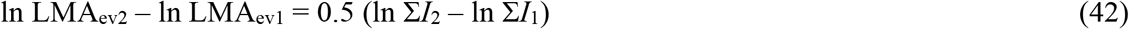

Similarly for the growing-season length and temperature:

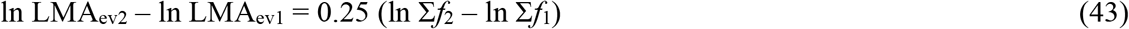

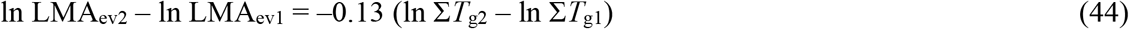

Thus, when moving from site 1 to site 2, the predicted changes in LMA are 50%, 25% and – 13% of the changes in PPFD, growing-season length and temperature respectively.

#### Environmental dependencies of LMA for deciduous species

As the lifespan of deciduous leaves is constrained by growing-season length, LL_de_ should also be proportional to *f* (LL_de_ = 365 *f*). Re-arranging equation (32) and imposing this additional constraint yields the following expression for deciduous LMA as a function of environment alone:

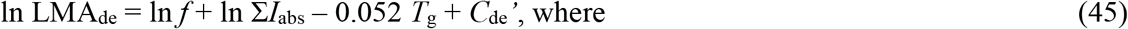

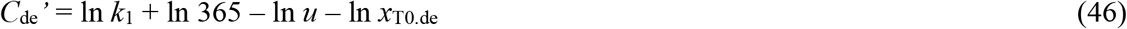

Equation (45) implies that deciduous LMA, unlike evergreen LMA, should increase in direct proportion to both PPFD and growing-season length, and decline with warming four times faster than in evergreens.

#### The aridity constraint on LMA

Finally, we account for the expected increase in LMA with aridity as an additional, physical constraint on LMA for both deciduous and evergreen species. This additional constraint could in principle have been included in equations (34) and (45) as an independent predictor. However, the current optimality framework does not predict the magnitude of this response. We therefore estimated the aridity effect empirically and used this additional information to modify the predicted within-site distribution of evergreen LMA, shifting the peak to higher values in drier sites.

#### Leaf trait data

We tested our quantitative predictions with the woody plants represented in two non-overlapping datasets: the GlopNet data set^1^, and a version of the China Plant Trait Database (CPTD)^5^ augmented with data compiled by Geng et al.^28^ from alpine shrublands. GlopNet contains paired LMA and LL data for 326 evergreen and 179 deciduous species at 45 sites, distributed globally. For deciduous plants we focus here on those whose growing season is constrained by a cold winter; thus, tropical deciduous species that shed leaves in the dry season have been excluded from the analysis. Applying this filtering criterion, the GlopNet data set contains data on LMA alone from 1225 evergreen and 98 deciduous species at 146 sites, distributed from 69°S to 40°N. The CPTD provides LMA data from 419 for evergreen and 398 deciduous species at 164 sites, spanning the range from alpine and boreal to tropical environments, and from deserts and steppes through woodlands and forests. Data in the CPTD were obtained using a defined sampling strategy to ensure adequate representation of major species in all strata^5^.

#### Site climate data

The Simple Process-Led Algorithms for Simulating Habitats (SPLASH) model was used to calculate site-specific bioclimatic variables from climatologies of monthly temperature, precipitation and sunshine hours^41^. Defining the (thermal) growing season as the period when mean quasi-daily temperature (interpolated from monthly data) is above 0°C, we calculated the ratio of growing-season length to the number of days in the year and the mean values of temperature and PPFD during growing season as estimates of site-mean *f, T*_*g*_ and PPFD. To account for the fact that the field-measured trait data reflect leaves developed at a range of irradiances at different levels in the canopy, we applied the approach used by Wang et al.^42^ to estimate *I*_abs_ from the site-mean PPFD, with the help of leaf area index values extracted from SeaWiFS data^43^. A moisture index, defined as the ratio of estimated annual actual to potential evapotranspiration (denoted α_p_), was also calculated using SPLASH. For the GlopNet data set, climatological data were extracted from Climate Research Unit data at a grid resolution of 10 minutes^44^. For the extended CPTD, climatological data were extracted from 1 km gridded data interpolated from 1814 meteorological stations (China Meteorological Administration, unpublished data: 740 stations have observations from 1971 to 2000, the rest from 1981 to 1990) using a three-dimensional thin-plate spline (ANUSPLIN version 4.36; Hancock and Hutchinson, 2006).

#### Statistical analysis

Ordinary Least-Squares (OLS) multiple regression was used to estimate the partial environmental dependencies of LL and LMA from data, for comparison with independent predictions from theory. All traits and bioclimatic variables except for temperature were log-transformed to facilitate comparisons with theoretical predictions, and to meet the assumptions of the OLS model. For consistency with our definition of the growing season we excluded deciduous species from sites with year-round temperatures > 0°C. These are presumed to be drought-deciduous. The effect of this exclusion could only be minor, however, as it merely reduces the sample size for deciduous LMA to 292 in GlopNet and 621 in the CPTD. Climatic controls of the relationship between LL and LMA were tested using the GlopNet paired LL and LMA data for evergreens. The constraint of growing-season length on LL in deciduous species was also tested using GlopNet data. The parameter *C*_*c*_ in equation (14) was estimated as the median of individually estimated values. Given the larger latitudinal range covered, GlopNet LMA data were used to investigate the opposing latitudinal trends in evergreen and deciduous species. Climatic controls on LMA were tested using both GlopNet and CPTD data, but CPTD-based results are given in the main text because of its larger sample size, especially for deciduous species. Stepwise regressions, with or without evergreen LL as a predictor, for GlopNet LMA data allowed us not only to test the explanatory power of LL alone on LMA, but also to test its indirect environmental effects on LMA when LL was excluded. For individual predictors that proved non-significant (probably due to the correlations between the bioclimatic variables), we further tested their effects on the residuals from the main model.

The prior distributions of LL and LMA were derived from GlopNet. We also used GlopNet to test the predicted within-site distributions of LMA. Six sites with adequate sample size were selected for this test, including two sites each from evergreen-dominant vegetation in tropical rainforests, temperate forests and woodlands (for details see Table S3). As shown in Figure S2, the prior distribution is constrained by the site-specific optimal relationship of ln LL to ln LMA with the slope set to unity and the intercept being the sum of the estimated value of *C* and the climatic terms in equation (14). The posterior distributions of ln LMA were generated after further imposing the aridity effect. As this effect is not theoretically quantified we used the best-match prediction obtained from regression of ln LMA data against climatic variables in the CPTD (Table S1).

## Acknowledgements

This research was supported by the National Natural Science Foundation of China (no. 32022052, 31971495), National Key R&D Program of China (no. 2018YFA0605400) grants and the generosity of Eric and Wendy Schmidt by recommendation of the Schmidt Futures program. ICP acknowledges support from the ERC-funded project REALM (Re-inventing Ecosystem And Land-surface Models, grant number 787203) and a High End Foreign Expert award at Tsinghua University (GDW20191100161). This work is a contribution to the Imperial College initiative on Grand Challenges in Ecosystems and the Environment (ICP). IJW and ICP acknowledge support from the Australian Research Council (DP170103410).

## Author Contributions

HW and ICP developed the quantitative theory, building on previous work by KK and XTX. XTX also contributed to theory development. HW carried out the analyses and constructed the display items using datasets compiled by HW and IJW. ICP, IJW and SCQ contributed to the interpretation of results. HW and ICP wrote the first draft and IJW, KK, XTX, SCQ and NCS contributed to successive drafts.

## Notes

### Competing Interest Statement

The authors have declared no competing interest.

