## Supplemental Tables and Figures for "Leaf economics explained by optimality principles" (from another version of this preprint: version 1, 2021-02-08)

**Table S1 | Summary statistics for the biotic and abiotic dependencies of leaf longevity based on the GlopNet data set.** Ordinary Least Squares multiple linear regression was fitted for the GlopNet data set on leaf longevity (LL) against explanatory variables for evergreen species (upper panel) and deciduous species (lower panel). The explanatory variables comprise leaf mass per area (LMA, g dry matter $m^{-2}$), site-mean leaf absorbed Photosynthetic Photon Flux Density ($I$, mol photon $m^{-2}$ $day^{-1}$) and growing-season length as a fraction of the year ($f$, day $day^{-1}$). Explanatory and response variables were natural-log transformed. Theoretical values of slopes were derived from the optimality framework.

| Quantity | Theoretical value | Fitted coefficient | Confidence intervals | | *P* value | $r^2$ | df |
|---|---|---|---|---|---|---|---|
| *evergreen* | | | | | | | |
| LMA | 1 | 0.94 | 0.81 | 1.07 | < 2e–16 | 0.42 | 322 |
| *I* | –1 | –1.67 | –2.09 | –1.26 | 4.37e–14 | | |
| *f* | –0.5 | –0.21 | –0.47 | 0.04 | 0.0991 | | |
| Intercept | NA | 5.91 | 4.92 | 6.91 | 1.14e–06 | | |
| *deciduous* | | | | | | | |
| *f* | 1 | 0.79 | 0.63 | 0.94 | <2e-16 | 0.50 | 97 |
| Intercept | 5.90 | 5.34 | 5.23 | 5.43 | <2e-16 | | |

**Table S2 | Summary statistics for the environmental dependencies of leaf mass per area based on the China Plant Trait Database.** Ordinary Least Squares multiple linear regressions were fitted for the China Plant Trait Database data on leaf mass per area (LMA, g dry matter C $m^{-2}$) against significant explanatory variables for evergreen species (upper panel) and deciduous species (lower panel). The environmental explanatory variables comprise site-mean leaf absorbed Photosynthetic Photon Flux Density ($I$, mol photon $m^{-2}$ $day^{-1}$) during the growing season, growing-season mean temperature ($T_g$, ˚C), the ratio of actual to potential evapotranspiration ($\alpha_p$, dimensionless) and growing-season length as a fraction of the year ($f$, day $day^{-1}$). Explanatory and response variables were natural-log transformed, except for $T_g$. Theoretical values of slopes were derived from the optimality framework.

$^{*}f$ was not significant in the multiple regression of LMA, but had a significant effect on the model residuals.

| Quantity | Theoretical value | Fitted coefficient | Confidence intervals | | *P* value | $r^2$ | df |
|---|---|---|---|---|---|---|---|
| *evergreen* | | | | | | | |
| $I$ | 0.50 | 0.66 | 0.22 | 1.10 | 0.00344 | 0.13 | 612 |
| $T_g$ | –0.013 | –0.02 | –0.03 | –0.01 | 4.89e–5 | | |
| $\alpha_p$ | <0 | –0.25 | –0.47 | –0.02 | 0.03010 | | |
| Intercept | NA | 3.07 | 1.91 | 4.23 | 2.64e–7 | | |
| $f^{*}$ | 0.25 | 0.45 | 0.25 | 0.67 | 1.55e–5 | 0.03 | |
| *deciduous* | | | | | | | |
| $I$ | 1 | 1.21 | 0.88 | 1.55 | 1.39e-12 | 0.53 | 613 |
| $f$ | 1 | 0.78 | 0.52 | 1.04 | 5.63e-09 | | |
| $T_g$ | –0.052 | –0.06 | –0.08 | –0.04 | 3.56e-08 | | |
| $\alpha_p$ | <0 | –0.60 | –0.75 | –0.46 | 4.51e-15 | | |
| Intercept | NA | 1.70 | 0.80 | 2.60 | < 2e-16 | | |

**Table S3 | Detailed site information for Figure 3**

| Figure panel | Resource | Latitude (˚N) | Longitude (˚E) | Elevation (m) | PPFD (mol $m^{-2}$ $day^{-1}$) | Biome | Number of observations |
|---|---|---|---|---|---|---|---|
| Fig 3a | GlopNet | 35.2 | 140.15 | 300 | 9.97 | Temperate forests | 8 |
| Fig 3b | GlopNet | –33.58 | 151.29 | 39 | 12.91 | Temperate forests | 19 |
| Fig 3c | GlopNet | 2.98 | 102.30 | 100 | 11.34 | Tropical forests | 24 |
| Fig 3d | GlopNet | 1.93 | –67.05 | 120 | 10.78 | Tropical forests | 24 |
| Fig 3e | GlopNet | –32.97 | 146.15 | 233 | 16.13 | Woodlands | 24 |
| Fig 3f | GlopNet | –32.98 | 146.15 | 233 | 16.13 | Woodlands | 21 |

**Figure S1 | Probability distribution functions fitted to GlopNet data on leaf mass per area (LMA, g dry matter m$^{-2}$) and leaf lifetime (LL, day).** The data were natural log-transformed. Means and variances are shown for all species (black), evergreen species only (dark green) and deciduous species only (light green).

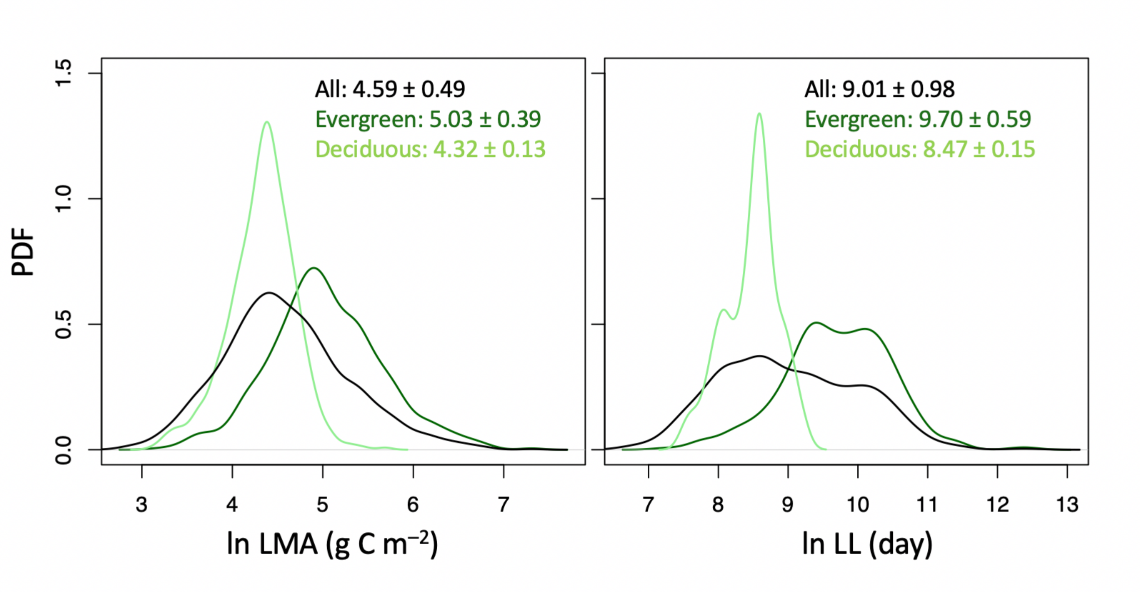

**Figure S2 | How the sensitivities of leaf mass per area (LMA) to environmental variables are determined by the prior distributions of LMA and leaf longevity (LL) and the site environment.** Solid green lines represent the combinations of ln LL and ln LMA that maximize average net carbon gain. The slopes of these lines are unity; the intercept is determined by climate (E1, E2). The green concentric circles represent the joint-normal distribution of ln LL and ln LMA and the resulting optimal LMA for evergreen species (green dots), which lie on the dashed green line, perpendicular to the solid green lines.

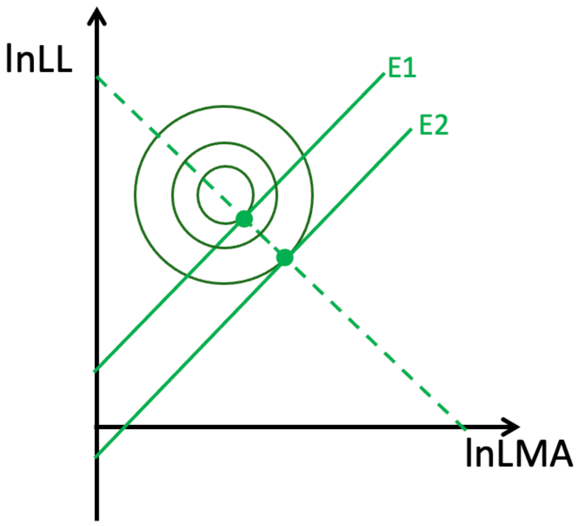

**Figure S3 | Climatic effects on the latitudinal trends of LMA.** (a) The residuals of the model of LMA. Panels (b), (d), (f) and (h) show predicted separate effects of moisture index, fractional growing season length, photosynthetic photon flux density and temperature, respectively. Panels (c), (e), (g) and (i) show the corresponding cumulative effects. Data and regression lines are in green for evergreen and orange for deciduous species. Dashed lines are theoretical predictions. Data from the GlopNet trait database.

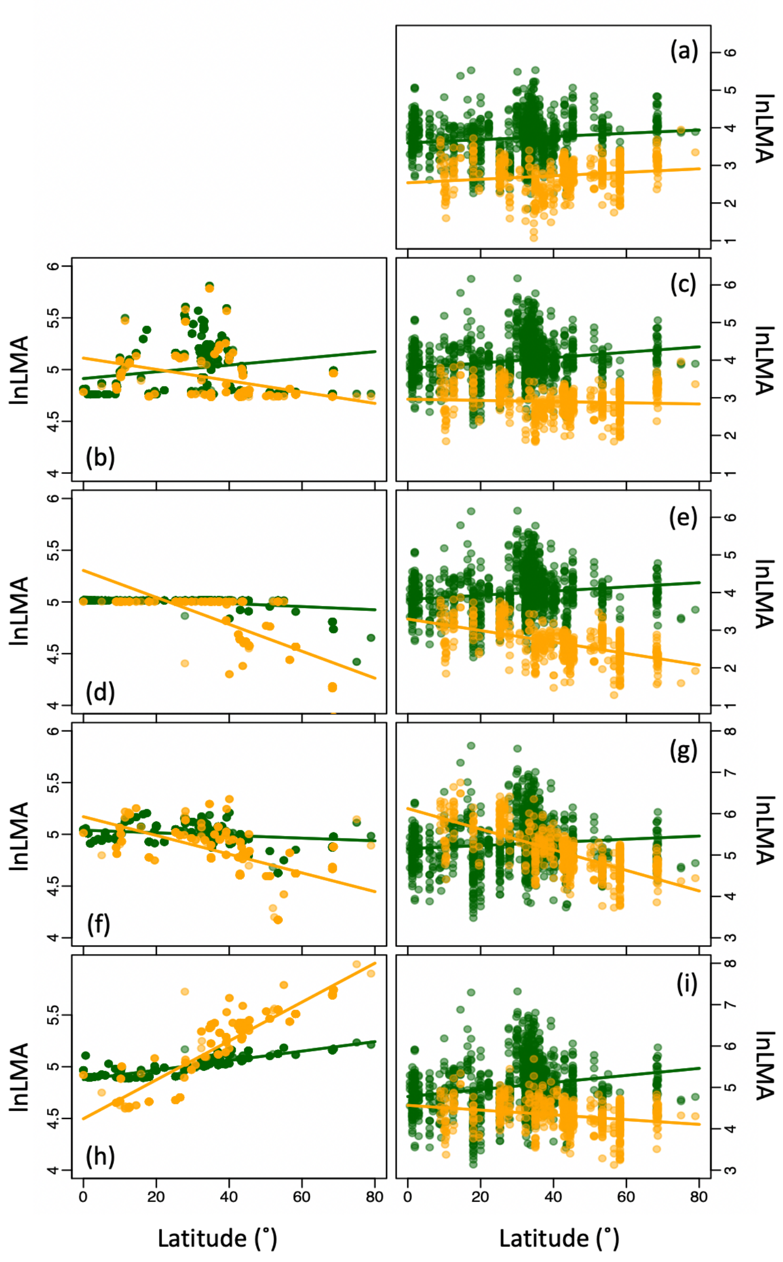
